# Epigenomics coverage data extraction and aggregation in R with tidyCoverage

**DOI:** 10.1101/2024.01.27.577537

**Authors:** Jacques Serizay, Romain Koszul

## Abstract

**Summary:** The *tidyCoverage* R package provides a framework for intuitive investigation of collections of genomic tracks over genomic features, relying on the principle of tidy data manipulation. It defines two data structures, *CoverageExperiment* and *AggregatedCoverage* classes, directly extending the *SummarizedExperiment* fundamental class, and introduces a principled approach to exploring genome-wide data. This infrastructure facilitates the extraction and manipulation of genomic coverage track data across individual or multiple sets of thousands of genomic loci. This allows the end user to rapidly visualize track coverage at individual genomic loci or aggregated coverage profiles over sets of genomic loci. *tidyCoverage* seamlessly combines with the existing Bioconductor ecosystem to accelerate the integration of genome-wide track data in epigenomic analysis workflows. *tidyCoverage* emerges as a valuable tool, contributing to the advancement of epigenomics research by promoting consistency, reproducibility, and accessibility in data analysis.

**Availability and implementation:** *tidyCoverage* is an R package freely available from Bioconductor ≥ 3.19 (https://www.bioconductor.org/packages/tidyCoverage) for R ≥ 4.4. The software is distributed under the MIT License and is accompanied by example files and data.

**Supplementary information:** Additional documentation is available from https://js2264.github.io/tidyCoverage/ and https://js2264.github.io/tidyCoverage/articles/tidyCoverage.html.

## 1. Introduction

Genome-wide epigenomic assays provide powerful methods to profile chromatin composition, conformation and activity. Linear “coverage” tracks are one of the main output files obtained when processing sequencing data. These coverage tracks, generally stored as .*bigwig* files, are often inspected in genome interactive browsers (e.g. IGV) to visually appreciate local or global variations in the coverage of specific genomic assays. Another approach to investigate genomic tracks is to compute and plot the average profile of a genomic track over a set of genomic loci. This approach is very efficient to summarize and compare the coverage of chromatin modalities (e.g. protein binding profiles from ChIP-seq, transcription profiles from RNA-seq, chromatin accessibility from ATAC-seq, …) over hundreds and up to thousands of genomic features of interest. This can be used to accurately describe, both qualitatively and quantitatively, multi-omic genomic tracks summarized across multiple sets of genomic features.

To create such metaplots, a number of softwares already exist in a command-line interface – e.g. *deeptools* (Ramírez et al., 2016) – or as packages in R – e.g. *genomation* (Akalin et al., 2015), *ATACseqQC* (Ou et al., 2018) or *soGGI* (Dharmalingam, n.d.). However, these softwares (1) are not interconnected to existing bioinformatic resources, (2) do not efficiently leverage the Bioconductor ecosystem and (3) do not use a tidy, intuitive syntax for data processing (Hutchison et al., 2023; Wickham et al., 2019). Here, we present *tidyCoverage*, an R package extending Bioconductor fundamental data structures and reusing principles of tidy data manipulation to extract and aggregate coverage tracks over multiple sets of genomic features.

## 2. Implementation

### 2.1. Two new S4 classes implemented from *SummarizedExperiment*

*tidyCoverage* implements the *CoverageExperiment* and *AggregatedCoverage* classes, both of which are built on top of the *SummarizedExperiment* class (Figure 1A). This ensures seamless creation and manipulation of these objects by end users, in particular those already familiar with popular packages built on top of *SummarizedExperiment*, such as *DESeq2* (Love et al., 2014) and *SingleCellExperiment (Amezquita et al., 2020). CoverageExperiment* objects organize a collection of genome-wide tracks (from local .*bigwig* files or numerical tracks stored in memory) and a collection of sets of genomic features of interest. When instantiated, the coverage of each genomic track over genomic features is extracted using the efficient underlying Bioconductor parallelization and import infrastructure (Lawrence et al., 2009) and stored in memory as an array. *AggregatedCoverage* further computes statistical metrics (e.g. mean, median, standard deviation, etc.) from the coverage vectors stored in a *CoverageExperiment* object. *tidyCoverage* data structures are natively compatible with other genomic data representations (e.g. *GenomicRanges, RleList, OrgDb*) and facilitate the integration of epigenomic data into large-scale multi-omics projects.

**Figure 1:**
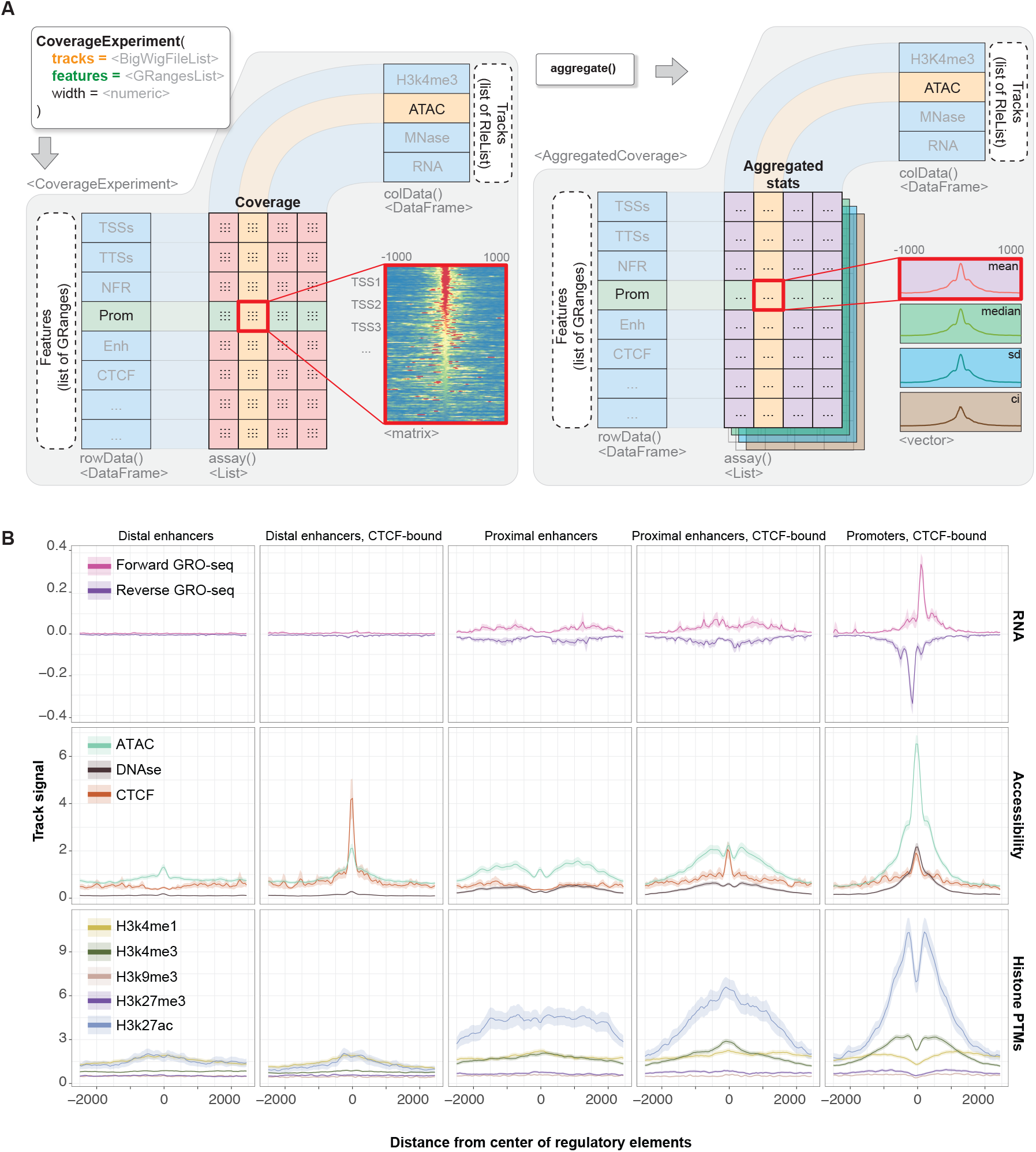
Overview of *tidyCoverage* functionalities. A. The *CoverageExperiment* object extracts and stores a separate coverage matrix for pairs of genomic track and genomic features. It can be further aggregated into a *AggregatedCoverage* object, which stores statistical metrics (mean, min, max, median, standard deviation, confidence interval) of the coverage of each track over each set of genomic features. B. *tidyCoverage* can be leveraged in combination with *ggplot2* functionalities to produce advanced aggregated coverage plots, for multiple tracks and genomic features.

### 2.2. Tidy principles for epigenomics

Tidy analysis of omics data has recently gained traction in large communities of bioinformaticians and programming languages (Hutchison et al., 2023), and *tidyCoverage* fully adheres to the tidy data paradigm. The package supports operative verbs defined in the tidyverse, such as *filter, mutate, group_by* or *expand* for *CoverageExperiment* and *AggregatedCoverage* objects. This enables researchers to efficiently organize, manipulate, and visualize epigenomic datasets in a tidy and structured format. *tidyCoverage* streamlines the intuitive exploration of large epigenomics datasets and facilitates data visualization using robust tools such as *ggplot2*.

## 3. Case studies

To demonstrate the usability of *tidyCoverage* package, we recovered 10 different epigenomic profiles in the human cell line GM12878 from the ENCODE data portal (Luo et al., 2020). We used *tidyCoverage* to extract track coverage over tens of thousands of ENCODE-annotated cis-regulatory elements, including promoters, proximal and distal enhancers (± CTCF). Aggregating epigenomics coverage highlighted the different composition, structure and activity of the chromatin which makes up different types of regulatory elements (Figure 1B). For instance, this reveals that CTCF enrichment is greater at distal enhancers than at proximal enhancers or promoters. This raises hypotheses regarding the implication of CTCF for chromatin looping and spatial folding at these different classes of regulatory elements.

## 4. Discussion

Compared to existing solutions, *tidyCoverage* focuses on data recovery and manipulation, using a standard representation of the data and principles of tidy data manipulation. *tidyCoverage* also ensures seamless integration of genomic track data into the existing genomics-centric Bioconductor ecosystem. This will contribute to the advancement of epigenomics research by fostering efficient and reproducible analyses.

## Code availability

The entire code used to generate Figure 1B is available here: https://github.com/js2264/tidyCoverage/blob/devel/vignettes/manuscript_figure.qmd.

## Data availability

All data presented in this manuscript have already been published. Human ENCODE-annotated regulatory elements were retrieved from (ENCODE Project Consortium et al., 2020) (Supplementary Table 10). The genomic tracks were retrieved from the ENCODE data portal from the following IDs: forward GRO-seq: ENCFF896TNM; reverse GRO-seq: ENCFF764SVR; Pol2RA ChIP-seq: ENCFF890SYC; CTCF ChIP-seq: ENCFF484SOD; DNAse-seq: ENCFF428XFI; ATAC-seq: ENCFF165WGA; H3K4me1 ChIP-seq: ENCFF785YET; H3K4me3 ChIP-seq: ENCFF736DCK; H3K9me3 ChIP-seq: ENCFF698SKV; H3K27me3 ChIP-seq: ENCFF119CAV; H3K27ac ChIP-seq: ENCFF458CR.

## Acknowledgements

We thank all our colleagues from the laboratory Régulation spatiale des génomes for fruitful discussions. This work was supported by fundings from the European Research Council under the Horizon 2020 Program grant agreement [771813], Q-life program and the Agence Nationale pour la Recherche to RK [ANR-22-CE12-0013-01, ANR-19-CE13-0027-02]. JS is recipient of a Postdoctoral fellowship from the Fondation ARC pour la recherche sur le cancer (ARC).

## Author contributions statement

**J**.**S**.: Conceptualization, Methodology, Software, Formal analysis, Investigation, Resources, Writing - Original Draft, Writing - other versions, Visualization

**R**.**K**.: Supervision, Project administration, Funding acquisition

## References

Akalin, A., Franke, V., Vlahoviček, K., Mason, C. E., & Schübeler, D. (2015). Genomation: a toolkit to summarize, annotate and visualize genomic intervals. Bioinformatics, 31(7), 1127–1129.

Amezquita, R. A., Lun, A. T. L., Becht, E., Carey, V. J., Carpp, L. N., Geistlinger, L., Marini, F., Rue-Albrecht, K., Risso, D., Soneson, C., Waldron, L., Pagès, H., Smith, M. L., Huber, W., Morgan, M., Gottardo, R., & Hicks, S. C. (2020). Orchestrating single-cell analysis with Bioconductor. Nature Methods, 17(2), 137–145.

Dharmalingam, G. (n.d.). Carroll T. soGGi: visualise ChIP-seq. MNase-Seq and Motif Occurrence as Aggregate Plots.

ENCODE Project Consortium, Moore, J. E., Purcaro, M. J., Pratt, H. E., Epstein, C. B., Shoresh, N., Adrian, J., Kawli, T., Davis, C. A., Dobin, A., Kaul, R., Halow, J., Van Nostrand, E. L., Freese, P., Gorkin, D. U., Shen, Y., He, Y., Mackiewicz, M., Pauli-Behn, F., … Weng, Z. (2020). Expanded encyclopaedias of DNA elements in the human and mouse genomes. Nature, 583(7818), 699–710.

Hutchison, W. J., Keyes, T. J., The Tidyomics Consortium, Crowell, L. H., Soneson, C., Yuan, V., Nahid, A. A., Mu, W., Park, J., Davis, E. S., Tang, M., Axisa, P. P., Kitt, J. W., Poon, C. L., Kosmac, M., Serizay, J., Sato, N., Gottardo, R., Morgan, M., … Mangiola, S. (2023). The tidyomics ecosystem: Enhancing omic data analyses. In bioRxiv (p. 2023.09.10.557072). 10.1101/2023.09.10.557072

Lawrence, M., Gentleman, R., & Carey, V. (2009). rtracklayer: an R package for interfacing with genome browsers. In Bioinformatics (Vol. 25, pp. 1841–1842). 10.1093/bioinformatics/btp328

Love, M. I., Huber, W., & Anders, S. (2014). Moderated estimation of fold change and dispersion for RNA-seq data with DESeq2. Genome Biology, 15(12), 550.

Luo, Y., Hitz, B. C., Gabdank, I., Hilton, J. A., Kagda, M. S., Lam, B., Myers, Z., Sud, P., Jou, J., Lin, K., Baymuradov, U. K., Graham, K., Litton, C., Miyasato, S. R., Strattan, J. S., Jolanki, O., Lee, J.-W., Tanaka, F. Y., Adenekan, P., … Cherry, J. M. (2020). New developments on the Encyclopedia of DNA Elements (ENCODE) data portal. Nucleic Acids Research, 48(D1), D882–D889.

Ou, J., Liu, H., Yu, J., Kelliher, M. A., Castilla, L. H., Lawson, N. D., & Zhu, L. J. (2018). ATACseqQC: a Bioconductor package for post-alignment quality assessment of ATAC-seq data. BMC Genomics, 19(1), 169.

Ramírez, F., Ryan, D. P., Grüning, B., Bhardwaj, V., Kilpert, F., Richter, A. S., Heyne, S., Dündar, F., & Manke, T. (2016). deepTools2: a next generation web server for deep-sequencing data analysis. Nucleic Acids Research, 44(W1), W160–W165.

Wickham, H., Averick, M., Bryan, J., Chang, W., McGowan, L., François, R., Grolemund, G., Hayes, A., Henry, L., Hester, J., Kuhn, M., Pedersen, T., Miller, E., Bache, S., Müller, K., Ooms, J., Robinson, D., Seidel, D., Spinu, V., … Yutani, H. (2019). Welcome to the Tidyverse. Journal of Open Source Software, 4(43), 1686.

